# Architecture of the type II secretion system

**DOI:** 10.64898/2026.05.07.723501

**Authors:** Minoosadat Tayebinia, Nouran Ghanem, Hui Zhang, Yu-Yuan Yang, Arianna Fornili, Vladimir Shevchik, Vidya Darbari, Richard Pickersgill

## Abstract

The virulence of emerging Gram-negative pathogens frequently arises from toxins delivered by the type II secretion system^1^. Cryo-EM single particle analysis and cryo-electron tomography and have defined the outer membrane secretin pore in detail, but the organisation of proteins within the periplasm and inner membrane that form the pilus assembly platform is not well resolved^2,3^. Here we combine AlphaFold^4^ models with single particle cryo-EM to define the organisation of the pilus assembly platform. We show that CLM heterotrimers form a continuous link from the cytoplasmic ATPase, across the inner membrane and periplasm, to the base of the secretin channel. AlphaFold models of the inner membrane spanning rotor and cytoplasmic ATPase fit readily within the cryo-EM density. The resolved secretion system exhibits an offset between the inner membrane assembly platform and the outer membrane secretin pore, together with profound asymmetry and an unexpectedly open periplasmic architecture. This architecture provides a route by which large, folded proteins access the secretion channel from the periplasm and suggests that substrate engagement may trigger the final steps in secretion system assembly leading to secretion.

## Main

The type II secretion system (T2SS) is a multiprotein complex that spans the cell envelope of many Gram-negative bacteria^5,6^. It secretes folded proteins, a toxin in the human pathogen *E. coli* IHE 3034 from the periplasm into the environment or onto the cell surface. The T2SS is a member of a superfamily of type IV filament assembling machines that build the type 4 pilus^7^ and several other related filaments^1,8–11^. These filament assembling machines have evolved from an ancient common ancestor and are fundamental to both bacterial and archaeal kingdoms^8^. The T2SS is a key virulence factor in many human pathogens including *Acinetobacter baumannii*^9^, *Klebsiella pneumoniae*^10^, *Pseudomonas aeruginosa*^11^ and *E. coli*^1^. It is discussed as an important drug target to prevent human disease^12^. Here we study the T2SS from enterotoxigenic *E. coli* IHE3034 which secretes the toxin SslE which causes meningitis and sepsis^13^. SslE is a 165kDa lipoprotein found in many pathogenic strains of *E. coli* which facilitates bacterial translocation through the mucosal barrier to get access to host cells^14^. The *E. coli* IHE3034 type II secretion system is encoded by 13 unidirectional genes *gspO, gspS* followed by 11 sequentially labelled genes *gspC* through *gspM* arranged in a single operon (Fig. 1A). This is the essential set of genes needed to assemble a functional type II secretion system; it is preceded by *sslE* gene for the secreted toxin. Here we refer to the *E. coli* T2SS proteins with the corresponding letters, C to S. The structure of the outer membrane secretin-pilotin complex comprising 15 copies of each of secretin D and pilotin S has been reported for several T2SS^2,15–18^ (Fig. 1B, C). There are two types of outer membrane secretin complex the *Klebsiella*-type and the *Vibrio*-type; the outer membrane secretin complex of *E. coli* IHE3034 is of the *Vibrio-*type^19^.

**Figure 1.**
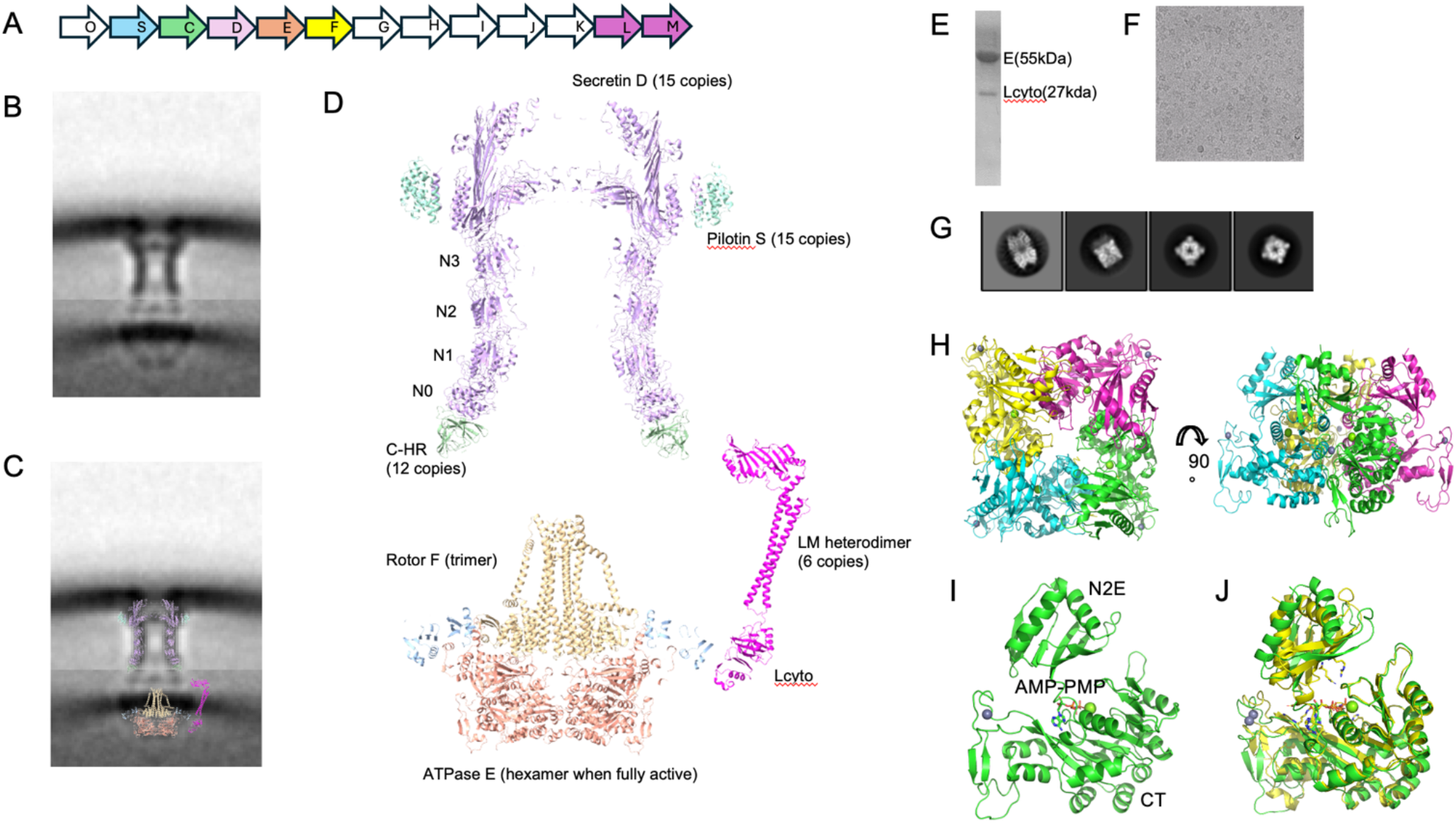
The type II secretion system is a multiprotein complex that translocates selected folded proteins through the outer membrane of Gram-negative bacteria. A) The operon for the T2SS from *E. coli* IHE3034 studied in this work. B) Visualization of the *L. pneumophilia* T2SS using electron cryotomography and subtomogram averaging. This panel was produced using Fiji^36^ and the EMDB density maps EMD-20712 and EMD-20713 for the outer and inner membrane components, respectively. C) The same composite map as shown in panel B) but with the *Klebsiella* secretin-pilotin complex (light magenta and sky blue) fitted to the outer membrane density and the AlphaFold model the EFLcyto complex (orange, yellow, blue; more details below) fitted to the inner membrane density. The AlphaFold model of the LM heterodimer is also shown (in magenta) displaced to the right-hand side of the EF complex to illustrate how the periplasmic ferredoxin-like domains might fit to the lobes of density between the inner and outer membrane components in the cryotomography map. D) A larger image of the molecules shown in panel C); the cross-section of the *Klebsiella* secretin and pilotin complex which comprises 15 copies of each of secretin D and pilotin S. A cross section of the secretin is shown in magenta and of the pilotin in sky blue. Also shown at the base of the secretin is the homology region of the inner membrane C protein (HR) in light green. The AlphaFold model of ATPase E is shown in orange, the rotor F in yellow, and the cytoplasmic region of the inner membrane protein L is shown in blue. The stoichiometry of the inner membrane assembly platform proteins as determined by Chenyatina and Low^2^ is 12C:6E:6L:6M. F is most plausibly a homotrimer^21^. The AlphaFold model of 6E:6Lcyto:3F captures the interaction of the N-terminal domain of E (N1E) and the cytoplasmic domain of L seen in the crystal structure of the individual domains. The structure of the secretin captures the known interaction of the HR domain of C (HR) and the N-terminal domain of the secretin D (N0) as seen in the crystal structure of the isolated domains. The L heterodimer structure captures correctly the known stoichiometry and interactions of the ferredoxin-like domains and the coiled-coil regions of L and M. E) – J). The structure of the IHE3034 ATPase. E) The cytoplasmic ATPase was purified in complex with the cytoplasmic domain of L (Lcyto) from the inner membrane assembly complex. F) High quality images of vitrified samples were recorded using cryo-EM. G) 2D-class averages reveal a back-to-back arrangement of E tetramers. H) The structure resolved at 3.3 Å reveals a compact organisation of the ATP subunits within each tetramer. I) The molecular details of the *E. coli* IHE3034 ATPase in complex with non-hydrolysable ATP (AMP-PNP) and magnesium (green sphere). The catalytic base Glu 328 on the CT domain is positioned for catalysis but arginine residues 204 and 219 (arginine finger residues) of the N2E domain are remote from the gamma-phosphate of the ATP analogue so this is an inactive arrangement of the N2E and CT domains. A zinc ion is shown as grey sphere binding to the CT domain. J) The orientation between the N2E and CT domains is different in this structure from other structures and from that seen in the forced hexamer 4KSS. This structure underlines the importance of the inner membrane assembly platform in organising a fully active hexameric ATPase. PyMOL was used to produce the molecular cartoons shown.

The inner membrane assembly platform comprises C, F, L and M plus the associated cytoplasmic ATPase E (Fig. 1D). It also includes the major pseudopilin G, and the minor pseudopilins H, I, J and K which are cleaved to their mature form by the inner membrane prepilin peptidase, O. Secretion of the substrate is driven by the motor E and inner membrane rotor of F. The ATPase E is anticipated to be hexameric when fully active and rotor F trimeric^20,21^; together they assemble the pilin subunits G into a helical pseudo-pilus which pushes the substrate into the environment^22^. The inner membrane protein C comprises a short N-terminal cytoplasmic region, a transmembrane helix followed by a long linker, the periplasmic HR domain and the C-terminal PDZ domain. The periplasmic HR domain interacts with the N-terminal secretin domain^23–25^ and the PDZ domain is implicated in substrate specificity^26^. L comprises a cytoplasmic domain that binds to the N1E domain of ATPase E^27^, a transmembrane helix followed by a long linker, and a periplasmic ferredoxin-like domain. The structure of M is like that of L but lacks an extended cytoplasmic region. The ferredoxin-like domains of L and M tend to form heterodimers suggesting that full-length LM is also likely to be a heterodimer^28^. A recent publication provides evidence that C, L and M form an assembly platform subcomplex XpsCLM from *Xanthomonas euvesicatoria*^*29*^.

### The inner membrane assembly complex directs assembly of the ATPase

We aimed to determine the structure of the *E. coli* IHE 3034 cytoplasmic ATPase in complex with the N-terminal cytoplasmic domain of the inner membrane protein L with which it interacts^27^. A non-hydrolysable ATP analogue (AMP-PMP) was used to stall the ATPase. The cryo-EM structure solved at 3.3 Å is a back-to-back dimer of tetramers with each protein subunit comprising the C-terminal domain CT and second N-terminal domain N2E domain of E (Fig. 1E-H; for image processing details see Supplementary Fig. 1). The N1E domain of E and the N-terminal cytoplasmic domain of L (Lcyto) are not seen in this structure implying a flexible arrangement of the N1E domain in the absence of the other inner membrane assembly platform proteins to restrain its orientation. An inactive arrangement of the CT and N2E domains is trapped in this structure because the arginine finger residues from the N2E domain are remote from the γ-phosphate of the ATP analogue bound to the CT domain (Fig. 1I). Remarkably despite the tightly packed tetrameric form the inter-molecular interface between the N2E and CT domains of adjacent chains is consistent with the other structures of this family of ATPases. The intra-molecular arrangement is different from previously seen and closest to the ADP-closed form in the forced hexamer of PilB (Fig. 1J). It is reasonable to conclude from this unusual tetrameric arrangement that the organisation of ATPase subunits in the fully assembled T2SS is orchestrated by the inner membrane assembly platform.

### The secretin directs assembly of the inner membrane assembly platform

The part of the T2SS operon comprising contiguous genes corresponding to the inner membrane assembly platform proteins, EFGHIJKLM (E-M), was expressed in *E. coli* C43 cells. The membrane proteins were solubilised using DDM. Purification using the Strep tag on E followed by size-exclusion chromatography gave three relatively pure protein bands corresponding to E, L and M (Fig. 2A); purification using a Flag Tag on F yielded four bands on SDS-PAGE corresponding to E, F, L and M (Fig. 2B). However, negatively stained samples of both these preparations showed considerable heterogeneity. There are many reasons why the inner membrane assembly platform may be highly heterogeneous in these samples and key among these is that the inner membrane assembly platform protein C which interacts with the outer membrane secretin D is missing in this construct. It is reasonable to conclude that the binding of the inner membrane protein C to the outer membrane secretin helps organise the architecture of the inner membrane assembly platform. Subsequent work therefore focused on purification of the full secretion system and subcomplexes from the full T2SS.

**Figure 2.**
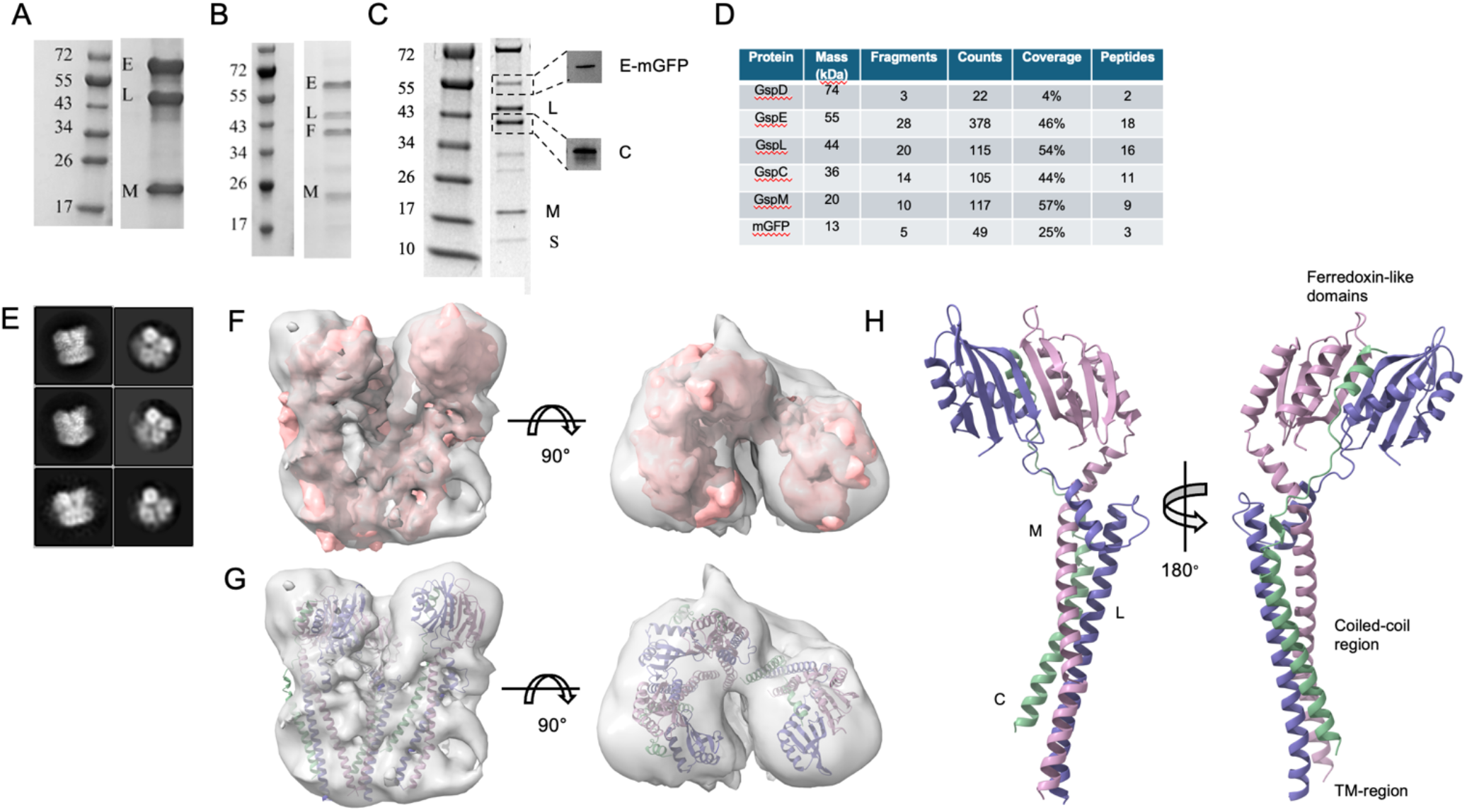
The structure of the CLM heterotrimer complex from the inner membrane assembly platform. A) The inner membrane complex (E-M) was expressed recombinantly and pulled out using the Strep tag on E. L and M are pulled out along with the ATPase E but the cryo-EM images showed highly heterogeneous particles (not shown). B) The inner membrane complex (E-M) was subsequently pulled out using the Flag tag on F. E, L and M are pulled out along with the rotor F but again the cryo-EM images showed highly heterogeneous particles. C) Purification of the *E. coli* IHE3034 T2SS (O-M) using a Flag-tag on the C-terminus of C. This protein also has a mini-GFP on the C-terminus of E. The Flag-tag on C pulls out C, D, L, M, S. D) Mass-spectrometry of the purified sample (shown in panel C) confirmed the presence of C, D, E, L, M. C:L:M are circa 1:1:1 in this sample. E) 2D-class averages from this sample are different from those of the secretin and the ATPase suggesting the presence of a different sub-complex. F) and G) the cryo-EM density together with three copies of the AlphaFold model of the CLM heterotrimer. H) The three CLM heterotrimers are the same comprising the heterodimer of L, M ferredoxin-like domains and the coiled-coil assembly of the alpha-helical regions of C, L and M. The HR and PDZ domains to the C-terminal region of C are not seen in the density presumably because they are flexible or disordered. Panels F) and G) were prepared using ChimeraX and panel H) using PyMOL.

### The CLM heterotrimer of the inner membrane assembly platform

The cloned IHE3034 T2SS operon producing proteins OSCDEFGHIJKLM (O-M; Fig 1A) was over-expressed in *E. coli* and purification of complexes exploited a construct engineered to have affinity tags on the N-terminus of E (Strep-tag) and C-terminus of C (Flag-tag). This recombinant T2SS secreted the toxin SslE or assembled a pilus when the toxin or major pilin protein were overexpressed together with the secretion system (Supplementary Fig.2). Assessing pilus formation by the recombinant T2SS was inspired by previous studies showing the T2SS could assemble pili^30^. If the construct expressing E-M was used in place of O-M, there was no secretion of SslE nor pilus generation confirming that the recombinant O-M IHE3034 secretion system is secretion competent but the E-M not. A mini-GFP was also introduced at the C-terminus of E in the construct used in this section, originally with the aim of facilitating correlated light and electron microscopy and cryo-electron tomography. Type II secretion system complexes were purified using the Flag-tag on C followed by density sedimentation in the presence of the cross-linker glutaldehyde (GraFix). The SDS-PAGE gel of the least dense fraction from the GraFix run shows the presence of C, D, E, L and M, a result confirmed by mass spectrometry (Fig. 2C, D). Particles imaged by negative stain could be segregated into those corresponding to the secretin-pilotin complex, those corresponding to ATPase and those corresponding to a third protein complex. The class averages for this third complex are shown in Fig. 2E. The initial maps showed strong anisotropy which was addressed by adding data collected using a series of tilt angles.

The cryo-EM density for this third complex at sub-nanometre resolution is consistent with three copies of the AlphaFold model of the CLM heterotrimer comprising associating ferredoxin-like domains of L and M and coiled-coil regions of L, M and C (Fig. 2F, G). The CLM heterotrimer was a slightly better fit to the density than the LM heterodimer. At this resolution the three CLM heterotrimers in this asymmetric inner membrane assembly platform subcomplex are essentially the same. The density reveals that the heterodimerisation of L and M ferredoxin-like domains evidenced by previous biochemical studies and crystal structures of the isolated domains^28,31–33^ occurs in the subcomplex isolated from the functional T2SS. Association of the coiled-coil regions of L, M, and C has been reported previously^33,34^. The AlphaFold model is consistent with the density map, and the transmembrane helices of C, L and M are in appropriate position for insertion into the inner membrane. The periplasmic HR and PDZ domains of C and cytoplasmic domain of L are not seen in the density and are presumed to be disordered in the structure (Fig. 2H). The CLM heterotrimer is consistent with the purification method, an affinity tag on C, and with mass-spectrometry of this sample which shows a stoichiometric ratio of C:L:M.

### The architecture of the *E. coli* IHE3034 type II secretion system

The cloned *E. coli* IHE3034 type II secretion system (O-M) was also produced with Strep-tag on E and Flag-tag on F. When purified using the Strep tag on E the SDS-PAGE shows the presence of six proteins: S, C, D, E, L, and M (Fig. 3A) whose identity were confirmed by mass-spectrometry (Fig. 3B). This purification was successful in pulling out the outer membrane secretin-pilotin complex (DS) as well as inner membrane proteins. This purified secretion system could be successfully imaged using negative stain, but the density of particles was low. The third and most successful purification method was by density sedimentation using the GraFix method alone. After sedimentation the least dense GraFix fraction was found to comprise the secretin-pilotin complex (Fig. 3C) and the densest fraction the full-length secretion system (Fig. 3D). The structure of the *E. coli* IHE3034 secretin-pilotin complex from the least dense GraFix fraction was refined at 3.85 Å resolution using C15 symmetry (Fig. 3E; for processing details see Supplementary Fig. 3). This structure is virtually identical to that previously solved from *E. coli* ETEC H10407^15^. When C15 symmetry is applied to refine the secretin complex from the heavy GraFix fraction where the inner membrane assembly platform is also present, then the average positions of the secretin N0 and inner membrane HR domain can be seen in density. The structure of the secretin-N0 domain/C-HR domain complex seen in the crystal structure^23^ fits into this averaged map (Fig. 3F). The weaker density for the N0 domains shows that these domains are less well ordered than the rest of the secretin. All 15 N0 domains are unlikely to be in complex with C-HR domains so the averaged density for the HR domains is expected to be weaker and this is what is seen.

**Figure 3.**
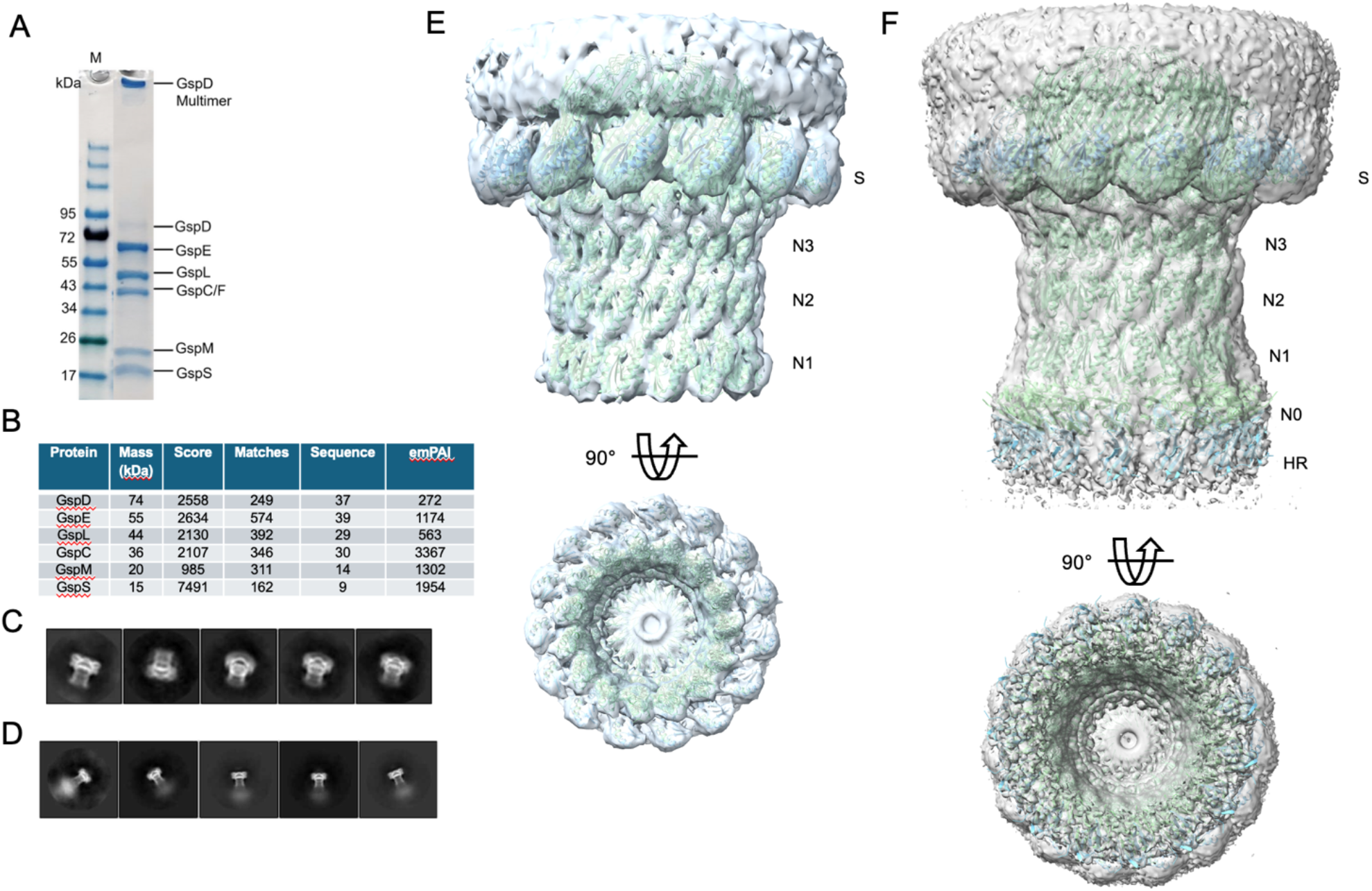
The structure of the secretin and secretin-pilotin complex. A) Purification of the full *E. coli* IHE3034 T2SS (O-M) using a Strep-tag on the N-terminus of E showing evidence for C, D, E, L, M, S. B) Confirmation of the presence of C, D, E, L, M, S from mass-spectrometry on bands cut from the gel. C) 2D-class averages showing that the least dense GraFix fraction corresponds to the secretin-pilotin complex. D) 2D-class averages showing that the densest GraFix fraction corresponds to the full-secretion system comprising well-ordered secretin and less well-defined inner membrane assembly platform. E) The structure of the secretin-pilotin complex solved using C15 symmetry from the least dense GraFix fraction (2D-class averages are in panel C). F) The structure of the secretin-pilotin complex solved using C15 symmetry from the densest GraFix fraction (2D-class averages are in panel D). There is density for the secretin N0 domain and the HR domain of the inner membrane protein C at the base of the secretin. The density supports a preferred orientation of the HR domain, but it is weaker which suggests flexibility, possibly multiple conformations, and lower occupancy of the HR domains compared to the secretin domains.

At this point to make further progress positioning the inner membrane proteins it was necessary to model in the absence of symmetry as the most remarkable characteristic of the inner membrane assembly platform in these samples is its asymmetry. The refined density in C1 for the full *E. coli* IHE3034 type II secretion system shows an offset inner membrane assembly platform of a volume into which the AlphaFold model of the cytoplasmic hexametric ATPase (E) and trimeric rotor protein (F) can fit (Fig. 4A; for processing details see Supplementary Fig. 4). The cryo-EM density is reassuringly similar in overall shape to that seen in the original lower resolution negatively stained micrograph of this complex (Fig. 4B). The C15 symmetry of the outer membrane secretin complex breaks down approaching the inner membrane assembly platform. In the cryo-EM density there are five density bridges between the inner and outer membrane parts of the complex. For four of these density bridges the density approaching the secretin are convincingly CLM heterotrimers (Fig. 4A) with the HR domains binding to the secretin N0 domains of secretin chains 1, 3, 5 and 12. To the inner membrane side of the CLM heterotrimers the cytoplasmic L domains are available to bind to the hexameric ATPase N1E domains as seen in the crystal structure of the isolated domains^27^. The position of the predicted transmembrane regions of C, L, M and F in this structure (C, L and M each have one transmembrane helix, and F has three transmembrane helices per subunit) are consistent with their positioning in the inner membrane (Fig. 4D). The fifth density bridge between inner membrane assembly platform and secretin is different and may correspond to a CLM heterotrimer in the process of associating with a proximal secretin domain. There is density for another three HR domains, but they are not associated with LM heterodimers so the density tethering them to the inner membrane is not seen. In total there are seven HR domains seen in an asymmetric distribution binding to the secretin N0 domains. Where there are no HR bridges to lock the ATPase and rotor to the secretin (secretin chains 6-9), the structure is more open, this leads to the offset position of the ATPase and rotor and large gaps in the architecture through which substrate could gain access. The observation of three HR domains bound to the secretin without adjacent LM heterodimers shows that individual C molecules also act alone as tethers between the inner membrane and outer membrane secretin as well as CLM heterotrimers. The complex has significant additional unmodelled density surrounding the modelled structure, some density appears to be less well-ordered LM heterodimers, or CLM heterotrimers, possibly also present are C, L and M monomers and pilin subunits. At the base of the secretin some additional density may be PDZ domains involved in substrate recruitment. Further improvements in the density and further corroborative experiments will be needed to assign these additional components. The detergent ring solubilising the transmembrane part of the secretin is clear in the density map, and it is likely that some of the additional density for the inner membrane platform is also detergent solubilising the inner membrane proteins.

**Figure 4.**
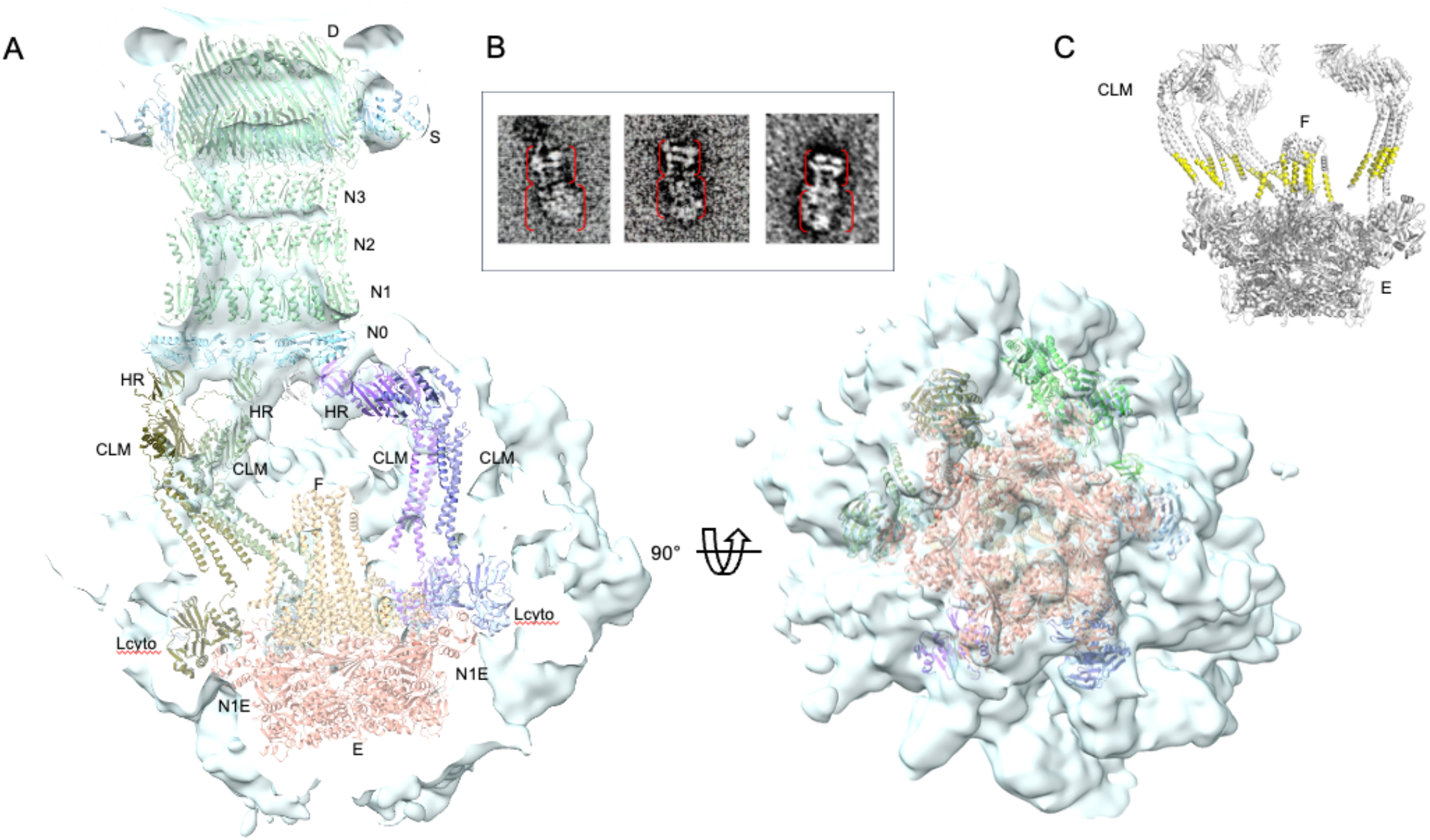
The structure of the full type II secretion system. A) A cross section of the cryo-EM density showing an asymmetric distribution of CLM heterotrimers around the base of the outer membrane secretin resulting in offset position of the ATPase E and rotor F. The CLM heterotrimers clip the secretin base to the ATPase. The HR domain of C binds to N0 domain of secretin D and the cytoplasmic domains of L (Lcyto) bind to the N-terminal N1E domain of the ATPase. This image captures the asymmetry and openness of the structure. B) The original negatively stained images of this complex showing the two distinct assemblies the outer membrane secretin and the inner membrane assembly platform. The inner membrane assembly platform is commonly offset in these negatively stained images. C) The transmembrane helices of C, L, M and F are highlighted in yellow to show that they are approximately in a plane consistent with their position in the inner membrane of the Gram-negative bacterial cell wall. Each F subunit has three transmembrane helices and each of the other proteins, C, L, M have a single transmembrane helix. Panels A) and C) were prepared using ChimeraX and PyMOL, respectively.

## Discussion

The structure resolved reveals the importance of the CLM heterotrimers in linking from the cytoplasmic ATPase through the inner membrane to the channel at the base of the secretin. The asymmetric distribution of the C-HR and LM ferredoxin-like domain heterodimers at the base of the secretin results in an offset between the inner membrane assembly platform and the outer membrane secretin pore. During this work we serendipitously resolved the well-ordered *E. coli* IHE3034 ArcAB-TolC multidrug efflux pump that spans inner and outer membranes^35^. The significance of this is that it confirms that the T2SS model presented has the correct spacing between inner membrane assembly platform and outer membrane secretin complexes. The remarkably open periplasmic architecture seen would allow for the recruitment of the large, folded protein substrate SslE to the secretion channel. It is plausible that substrate engagement triggers secretion by bringing the assembly platform into greater alignment with the secretin pore establishing six CLM links between secretin and the ATPase thereby stabilising the active E hexamer which then drives pilus assembly and secretion.

## Methods

### Cloning, protein expression and purification

The operon encoding the type II secretion system (*gspOSCDEFGHIJKLM*) was amplified from the genome of *E. coli* O18:K1:H7 IHE3034 and cloned into *pASK-IBA3c* to produce *gspO-M* with a Strep-tag at the N-terminus of E. Over 30 variants with affinity tags at different positions, mini-GFP, and cross-links between proteins were built on this background and four of these variants were used in this work (see supplementary Table 1). The variant that was used for the structure of the full T2SS and the secretin-pilotin complex had a Strep-tag on the N-terminus of E and a Flag-tag on the C-terminus F. The variant used to produce the CLM complex had the N-terminal Strep-tag at the N-terminus of E, a mini-GFP followed by a hexahistidine tag at the C-terminus of E and a Flag-tag at the C-terminus of C. These additional tags were added to the original construct using infusion, NEB’s Q5® Site-Directed Mutagenesis Kit or for the CLM complex structure was delivered by GenScript Biotech (UK). In another experiment, the cloned genes corresponding to the ATPase and the cytoplasmic domain of L were expressed leading to the ATPase tetramer structure. All plasmids were sequenced to ensure the sequences were correct. The clones were transformed into *E. coli* C43 (DE3) electro-competent cells (Lucigen) cells for protein expression using standard methods. Cells were grown on selective LB-agarose plates with chloramphenicol (30 μg/ml). Anhydrotetracycline (AHT, 0.2 mg/L) was added after the initial growth period to induce expression, and cells were grown for circa16 hours at 18°C. Cells were harvested by centrifugation at 5000 ×g for 20 minutes and pellets were re-suspended in ice-cold lysis buffer (20 mM Tris-HCl pH8.0, 150 mM NaCl, 1mM EDTA, 1mM DTT, DNase, 1mg/ml lysozyme and protein inhibitor cocktail tablets (Roche), followed by sonication on ice. The lysate was clarified by centrifugation at 18,000 × g for 30 min at 4°C. The membrane fraction was then collected by centrifugation at 140,000 × g for 1h at 4°C and resuspended in washing buffer (20 mM Tris-HCl pH8.0, 150 mM NaCl, 1 mM EDTA). The resuspended sample was centrifuged at 140000 ×g for an additional 30 minutes at 4°C. For the solubilisation step, the membrane pellet was resuspended in 20 mM Tris-HCl, pH 8.0, 150 mM NaCl,1 mM EDTA, 1% DDM (Anatrace), protease inhibitor cocktail, and 10% glycerol at room temperature for 1 hour. The solubilized sample was clarified by centrifugation at 132000 × g for 20 min at 4°C.

The solubilised membrane fraction containing the T2SS was applied directly for Gradient centrifugation or loaded on Strep-Tactin®XT 4Flow® resin (IBA Lifesciences). For the affinity purification Strep-Tactin®XT 4Flow® resin pre-equilibrated with 50 mM Tris-HCl (pH 7.6), 150 mM NaCl, 1 mM EDTA, 1 mM TCEP and 0.06% DDM. For elution, the same buffer was supplemented with 50mM Biotin. Flag-tag purification of the CLM complex used Anti-Flag M2 affinity gel (Sigma-Aldrich) washed 10x column volume with 50mM Tris-HCl pH 7.6, 150mM NaCl, 1mM EDTA PH 8, 0.06% DDM followed by 5x column volume with 50mM HEPES pH 7.6, 150mM NaCl, 1mM EDTA PH 8, 0.06% w/v DDM at 4 °C. Bound protein was eluted with 5x column volume of 100 µg/mL FLAG peptide (Sigma) in 50mM HEPES pH7.6, 150mM NaCl, 1mM EDTA PH 8, 0.06% w/v DDM. Peak fractions were pooled and cross linked with 0.1% glutaraldehyde (Sigma-EM grade) for 20 min on ice. Protein-containing fractions were analysed by SDS-PAGE and excised bands used for identification using LC-MS/MS.

GraFix gradients were prepared in Beckman Ultra-Clear 4.2 ml 11 × 60 mm ultracentrifugation tubes. Stepwise layers of 0.9 ml buffer comprising 50mM HEPES pH 7.6, 150mM NaCl, 1mM EDTA PH 8, 0.06% w/v DDM containing 40%v/v glycerol plus 0.1% glutaraldehyde, 30% glycerol plus 0.05% glutaraldehyde, 20% glycerol plus 0.025% glutaraldehyde, and 10% glycerol with no cross-linker were sequentially frozen in liquid nitrogen. A continuous gradient was formed by thawing at 4 °C. Cross-linked protein samples were layered on top and centrifuged in a Beckman SW60 Ti rotor at 25,000 rpm for circa15 hours at 4 °C. 150 µl fractions were collected using a syringe pump connected to an AKTA purifier. Peak-containing fractions were quenched with 100 mM Tris-HCl pH 7.5 and buffer exchanged using Amicon Ultra centrifugal filters 100KDa cut-off against 50mM HEPES pH 7.6, 150mM NaCl, 1mM EDTA PH 8, 0.06% w/v DDM to remove any traces of glycerol and Tris-HCl.

### Secretion competence assay

This method evaluates the capability of the recombinant T2SS to transfer its toxin SslE into the culture media. The *sslE* gene was amplified from *E. coli* IHE3034 chromosomal DNA and cloned into *pRSF*-Duet vector. The construct was co-transformed into *E. coli* C43 (DE3) along with *pASK3c*+*gspO-M*. Separately the *gspG* gene was used in place of the *sslE* construct to assess pilus formation. Maintenance and induction of the Duet plasmid used 50 μg/mL kanamycin and 0.5 mM IPTG, respectively. Maintenance of the *pASK3c* plasmid used chloramphenicol with induction using anhydrotetracycline as described above for the T2SS.

### Mass-spectrometry

Mass spectrometry was employed to identify the protein bands on SDS-PAGE or proteins present in the sample. The protein bands were excised from the gel and forwarded to the Mass Spectrometry unit at Cambridge Centre for Proteomics (Fig. 4B) or purified samples were analysed (Fig. 3D).

### Electron microscopy sample preparation and data collection

Negatively stained samples of the membrane protein complexes were prepared using ultra-thin support film (3 nm) on lacey carbon copper grids (Agar Scientific) glow discharged using the easiGlow system for 1 minute. 5 µl of protein was applied for 1 minute before blotting, then washed three times with 5 µl distilled water and subsequently stained with 5 µl of 1% uranyl acetate solution for 10 seconds before blotting again. These grids were examined using a JEM-1400 Flash microscope and images analysed using ImageJ.

For Cryo-EM structure determination of the T2SS the Leica EM GP2 Automatic Plunge Freezer with chamber at 4°C and 95% humidity was used. 4 μl of the complex was applied to glow discharged ultrathin carbon-coated Quantifoil R1.2/1.3 300 mesh Cu grids for 30 s before 4.5 s blotting time and vitrification in liquid ethane. Grids were assessed using the in-house 200 kV Jeol 2100 plus microscope with OneView 4K camera with SerialEM software. Data were collected using a Titan Krios (LonCEM & eBIC), with a total dose of 50 e/Å^2^. The pixel size was 0.829 Å for the secretin-pilotin sample and 1.08 Å for the full secretin system. Approximately 18,000 movies were collected from the secretin-pilotin sample, and 10,000 from the entire T2SS (densest GraFix sample). The CLM data were collected using a similar procedure and 0.84 Å pixel size.

The ATPase was incubated with 1mM AMP-PNP for 20 minutes on ice before applying 4 μl to glow discharged ultrathin carbon-coated Quantifoil R1.2/1.3 400 mesh Au grids which were incubated for 10s before blotting for 4.5 s and subsequent vitrification. Images were taken using a Titan Krios (LonCEM & eBIC) with Gatan K3 Summit camera in dose-fractionation mode at a calibrated magnification of 105k, corresponding to 0.85Å per physical pixel. The total dose was 40 e/Å^2^. Fully automated data collection used EPU with a nominal defocus range set from -1.5 to - 3.5 μm.

### Image processing

For the ATPase structure (Supplementary Figure 1) a total of 3613 movies were collected at super-resolution (0.425 Å/pixel). Data were processed using RELION3. Two rounds of 3D classification gave circa 269,000 particles (38.2%) which were used for 3D auto-refinement using C4-symmetry, followed by CTF refinement, Bayesian polishing and post-processing to give the final map at 3.25 Å resolution. All resolutions were estimated based on gold standard Fourier shell correlations (FSC)= 0.143 criterion.

For the CLM complex (Supplementary Figure 3) 840,245 particles were extracted with a 300-pixel box size and subsequently multiple rounds of 2D classification were used to remove low quality and heterogenous particles, resulting in a stack of 172,401 particles. These were used to generate a 3D initial model followed by 3D classification into three classes. Two classes containing 77,597 particles were selected for final refinement, yielding a reconstruction at 6.8 Å resolution. However, the map displayed strong anisotropy. To address this, two additional datasets were acquired at -20°, -30°, -35°, and -40° tilts. After motion correction and CTF estimation as above, 55,660 tilted micrographs were processed in RELION5^37^. Particles from the tilted datasets were cleaned by iterative 2D classification and merged with the refined particles from 0° tilt dataset, giving a combined total of 859,603 particles. Further 2D classification reduced these to 556,853 particles. Three classes of 3D initial model were generated and only one class containing 218,517 particles was used for 3D classification to obtain one class that was used for the final refinement. This construction reached 7.2 Å resolution and the unmasked map was significantly less anisotropic.

For the secretin and the full-length T2SS, collected movies were motion corrected using MotionCor2, and all subsequent image processing steps used CryoSPARC v4.7.1^38^. Contrast transfer function (CTF) estimation used Patch CTF Estimation. For the secretin dataset (Supplementary Figure 4), particles were initially picked by the general model with t = 0.15 in crYOLO. A total of 83,000 particles were picked and extracted with a box size of 600 pixels and 3× binning (pixel size 2.49 Å) for downstream processing and were used as input for 2D classification to remove the junk. Ab initio reconstruction (C1 symmetry) resulted in a flattened map showing strong preferred particle orientation in the dataset. To improve particle picking, the micrographs were denoised by Topaz and new templates were made from EMD-6917 for template picking, resulting in 70,874 particles. These were extracted with a 600-pixel box size (pixel size 0.829 Å) and passed through multiple rounds of 2D classification. The final 68,555 particles were refined using a 30 Å low-pass-filtered map (EMD-6917) as the initial volume, applying C15 symmetry and yielding a resolution of 2.7 Å, estimated using the Fourier shell correlation (FSC) 0.143 criterion.

For the full-length secretin system dataset (Supplementary Figure 5), a total of 260,000 micrographs were used for CTF estimation. Initial picking using crYOLO, followed by template-based picking in Topaz, resulted in 17,000 particles, after which particles were extracted with a box size of 800 pixels and 2× binning (pixel size 2.16 Å). The particles were 2D classified for dataset cleaning and were used to generate three initial volumes using the CryoSPARC *ab initio* reconstruction algorithm, which produced one class corresponding to the full-length secretin system. These volumes were then used as reference volumes to classify all extracted particles using heterogeneous refinement. A final 7,722 particles belonging to the full-length complex were refined using homogeneous refinement with C1 and C15 symmetry. This resulted in two maps, the asymmetric secretion system at a global resolution of 8.34 Å and secretin-pilotin-HR density at 4.37 Å resolution, as estimated by the Fourier shell correlation (FSC) 0.143 criterion. The density was further improved using LocScale-2.0^39^ to give the final map which is presented here and deposited.

### Model building and validation

The structure of the tetrameric ATPase was refined at circa 3.3 Å using COOT^40^ and Phenix^41,42^ starting from an initial model based on GspE from *Vibrio cholerae* (PDB code 4PHT). The structure of the CLM heterotrimer was the AlphaFold^4^ model refined using rigid-body optimisation of the coiled-coil regions and ferredoxin-like domains in Flex-EM^43,44^. The structure of the IHE3034 secretin-pilotin complex present in the light GraFix fraction was refined to at 3.85 Å resolution using iterative cycles of COOT or ChimeraX/ISOLDE^46,49^, and Phenix^50^, with C15 symmetry applied in Phenix. The initial model was based on the virtually identical secretin-pilotin structure determined from enterotoxigenic *E. coli*^*18*^ (PDB code: 5ZDH). C15 symmetry was applied to the heavy GraFix Fraction of the full T2SS and ChimeraX used together with an AlphaFold model of the N0/HR complex to produce the C15 secretin-pilotin-HR model. For the full secretion system, the AlphaFold models of 5 CLM heterotrimers and the AlphaFold model of the hexameric motor ATPase E in complex with trimer rotor F with the cytoplasmic domain of L bound to N1E of E, were fitted to the density using ChimeraX. The AlphaFold model of 6E6Lcyto3F was too large to generate in one run, so the three-fold symmetry of the complex was exploited in making the model. Additional HR domains where their identify was clear from their position were also fitted to the density. Flexible loop regions between fitted HR domains and the coiled coil regions of C were built using Boltz-2x^47^. The flexible loops between the coiled coil regions of L and Lcyto were built using the same process. Any chain clashes in composite AlphaFold models or between the 66 polypeptide chains fitted to the density were resolved using ChimeraX/ISOLDE.

## Author Contributions

MT molecular biology, biochemistry, electron microscopy; NG molecular biology, biochemistry, electron microscopy; HZ cloning of the *E. coil* IHE3034 secretion system; cryo-EM; YYY molecular modelling; AF supervision of molecular modelling; VD supervision of single particle analysis and single particle analysis; RWP fund raising, map interpretation, editing. All authors contributed to writing sections of the manuscript.

## Acknowledgements

This work was supported by the Biotechnology and Biological Sciences Research Council (BBSRC) Grant BB/W006693/1. Samples were screened using our in-house Jeol JEM 2100 plus microscope funded by BBSRC (BB/R000514/1). High-resolution data were collected at LonCEM using the Titan Krios cryo-EM or at the national electron Bio-Imaging Centre (eBIC; proposal BI35335). We are grateful to Nora Cronin for help with data collection at LonCEM. This research used Queen Mary’s Apocrita HPC facility, supported by QMUL Research-IT. We thank James Garnett for genomic DNA from *E. coli* IHE3034 from which we cloned the T2SS.

## Data availability

3D cryo-EM density maps produced in this work have been deposited in the Electron Microscopy Data Bank with accession code EMD-57257 (ATPase), EMD-57244 (CLM), EMD-57314 (Secretin), EMD-57315 (full T2SS). Atomic coordinates have been deposited in the Protein Data Bank (PDB) with accession code 29NU, 29KS, 29QZ and 29RA.

## Supplementary Material

Supplementary Figure 1. Cryo-EM workflow for the ATPase

Supplementary Figure 2. The recombinant T2SS is secretion competent

Supplementary Figure 3. Cryo-EM workflow for the CLM heterotrimer complex

Supplementary Figure 4. Cryo-EM workflow for secretin-pilotin complex

Supplementary Figure 5. Cryo-EM workflow for type II secretion system

Supplementary Table 1. Constructs used in this work

