## Supplementary Material for "Architecture of the type II secretion system"

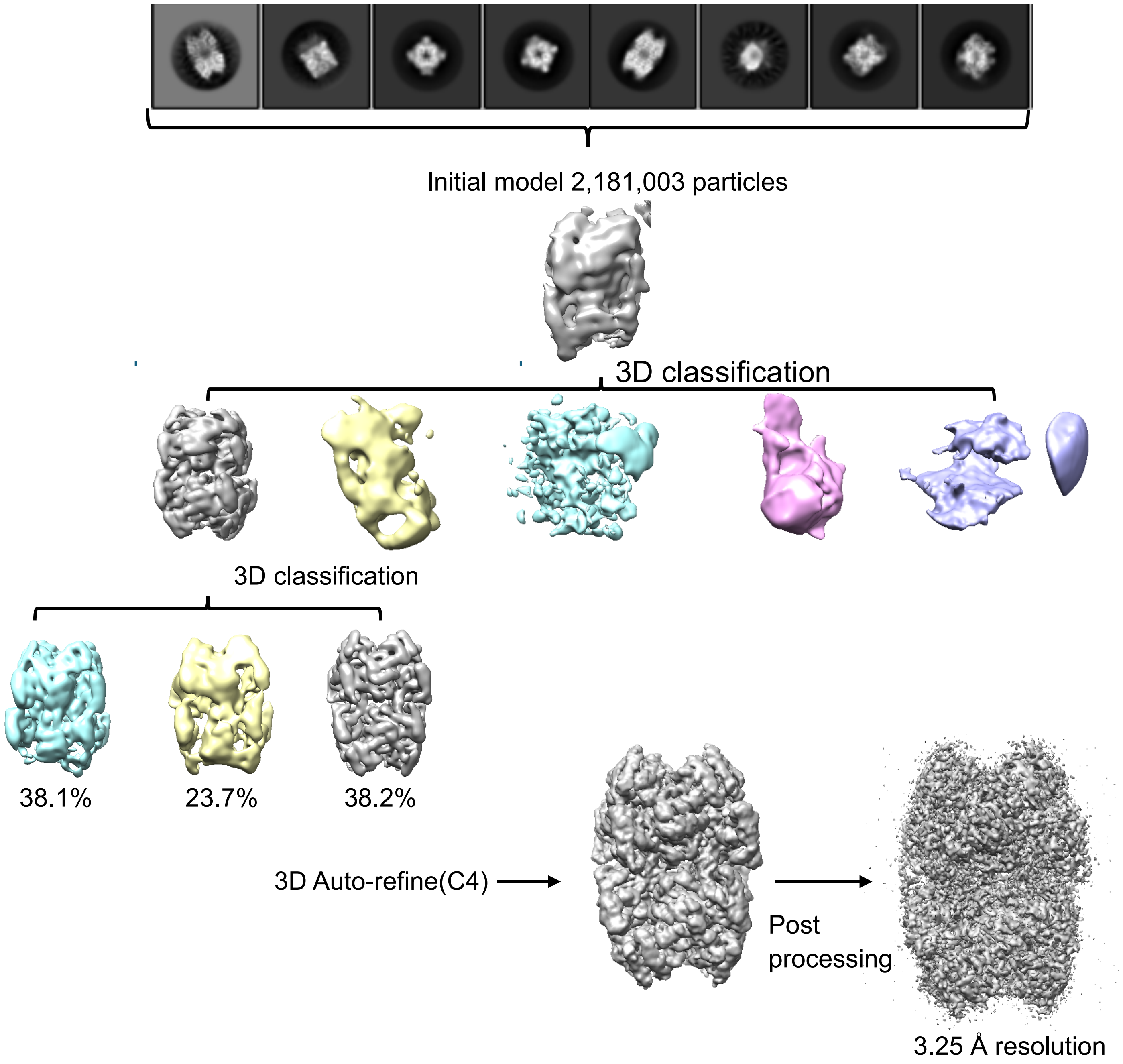


**Supplementary Figure 1. *E. coli* IHE3034 ATPase structure.** Cryo-EM single particle analysis workflow.

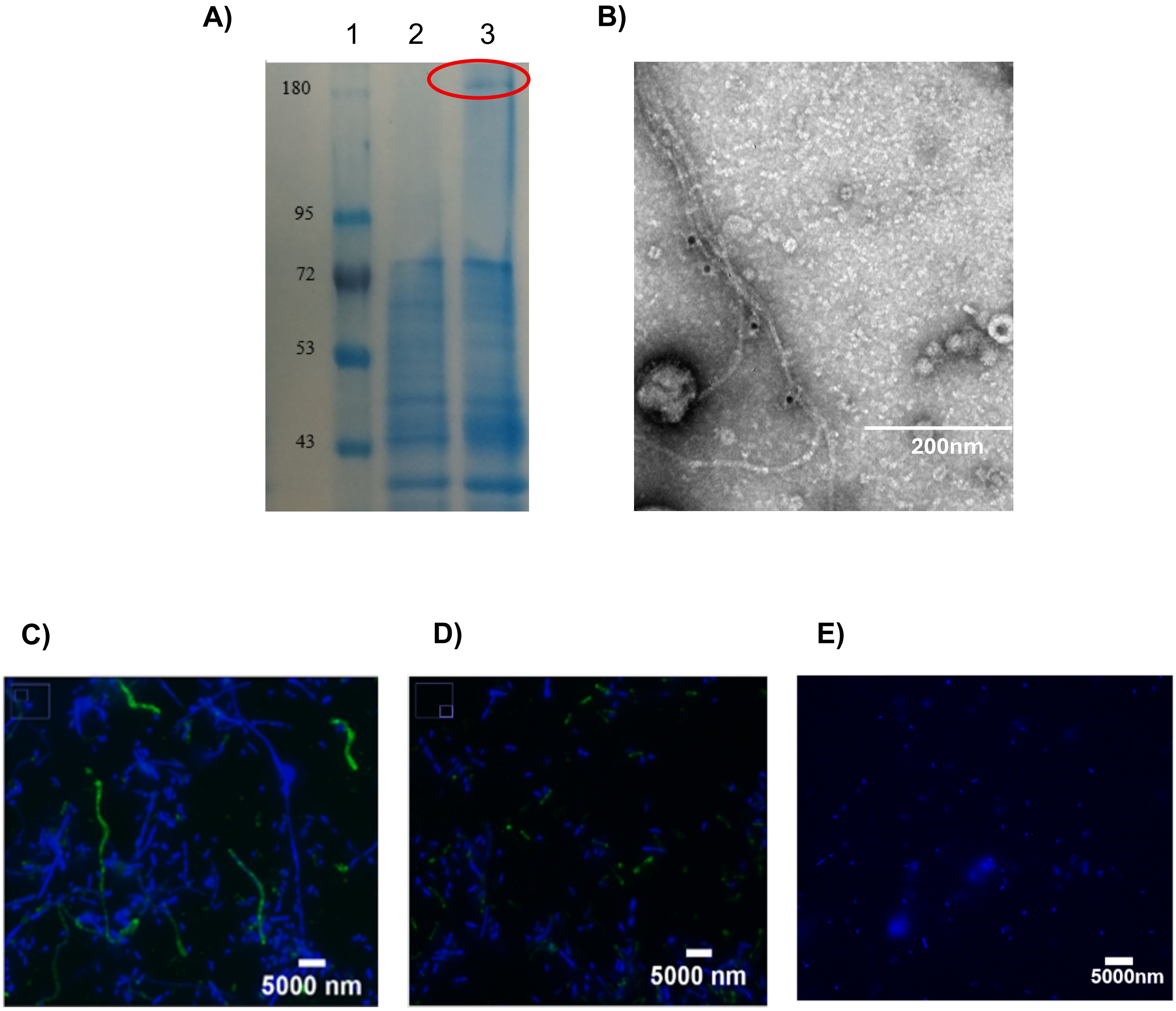


**Supplementary Figure 2.** The recombinant *E. coli* IHE3034 T2SS is secretion competent. **A**) SDS-PAGE gel. When the full T2SS (O-M) is expressed with the gene for SslE the toxin is secreted into the medium (lane 3, SslE is ringed in red). Lane 2 shows the control with the proteins corresponding to the partial T2SS (E-M) expressed, no SslE is secreted. Lane 1 has the molecular mass markers. **B**) Negatively stained image of the pili with immmunogold labelling of the Flag-tag on the major pilin subunit, GspG. The labelling was done on the grid which together with the burial of many of the Flag-tags accounts for the sparcity of labelling. Flag tags presumeably have limited accessibility in the pilus and the labvelling was done after the sample was applied to the grid. **C)** Overexpression of the major pilin subunit,

GspG, together with the T2SS which causes hyper-pili to be produced. The presence of green flourescently labelled hyperpili when the major pilin GspG is overexpressed along with the T2SS (O-M) is clear. **D)** No clear hyperpili when the major pilin subuint is overexpressed with the E-M construct co-expressing the contiguous inner membrane assembly proteins of the T2SS. **E**) No hyperpilus production when the active site glutamate of the ATPase (Fig. 1F) GspE was mutated to alanine. The pilus was detected using an anti-Flag antibody conjugated with DylLight 488 (shown in green on the micrograph). Nucleic acid was stained blue using DAPI.

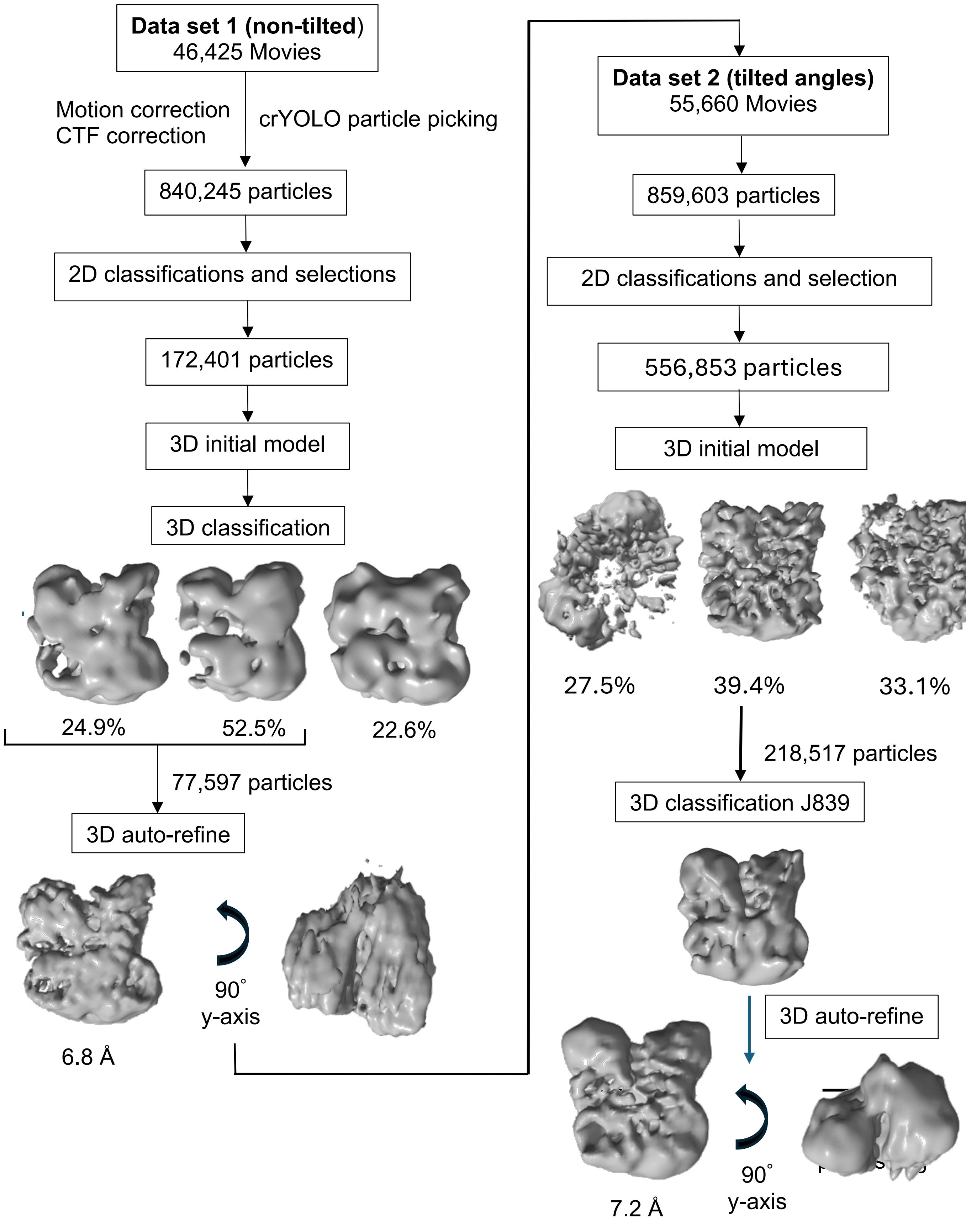


**Supplementary Figure 3. Cryo-EM single particle analysis workflow for the CLM inner membrane subcomplex.**

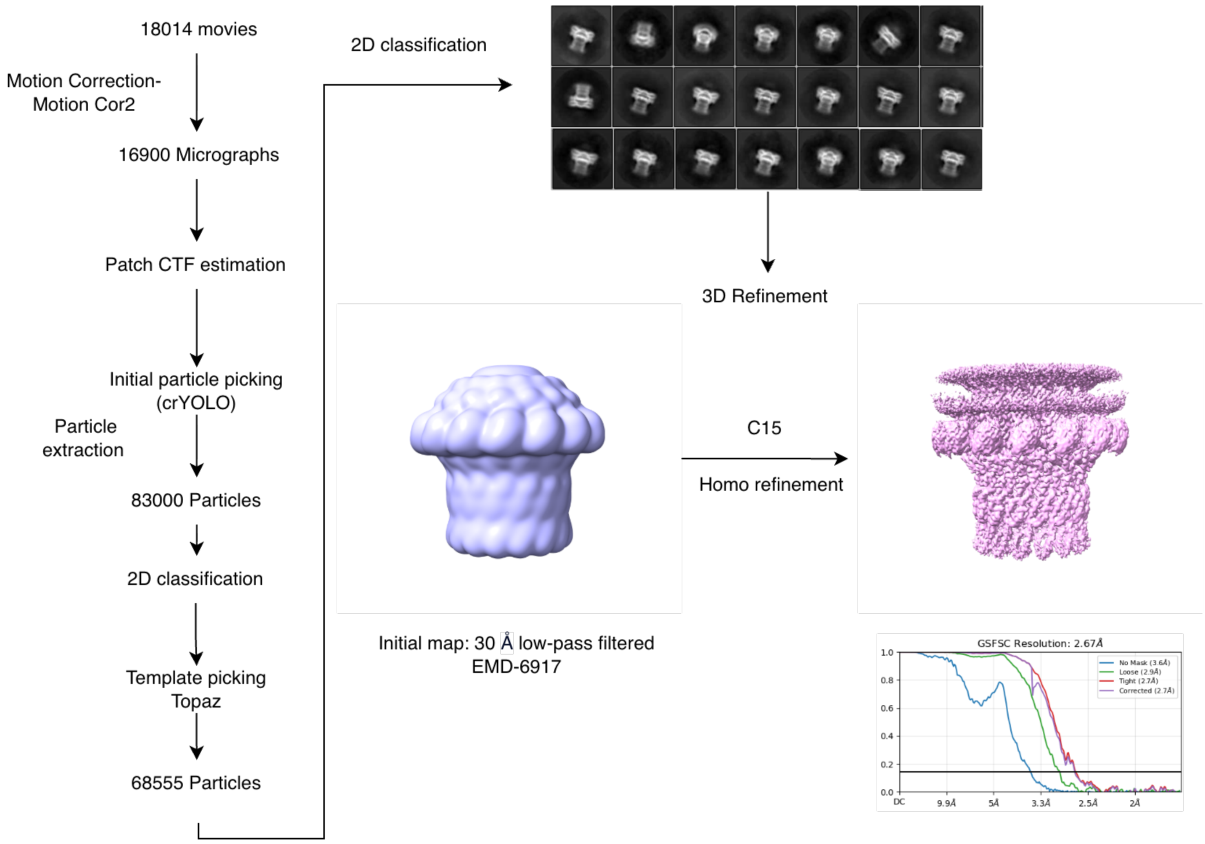


**Supplementary Figure 4. Cryo-EM single particle analysis workflow for the secretin-pilotin subcomplex**

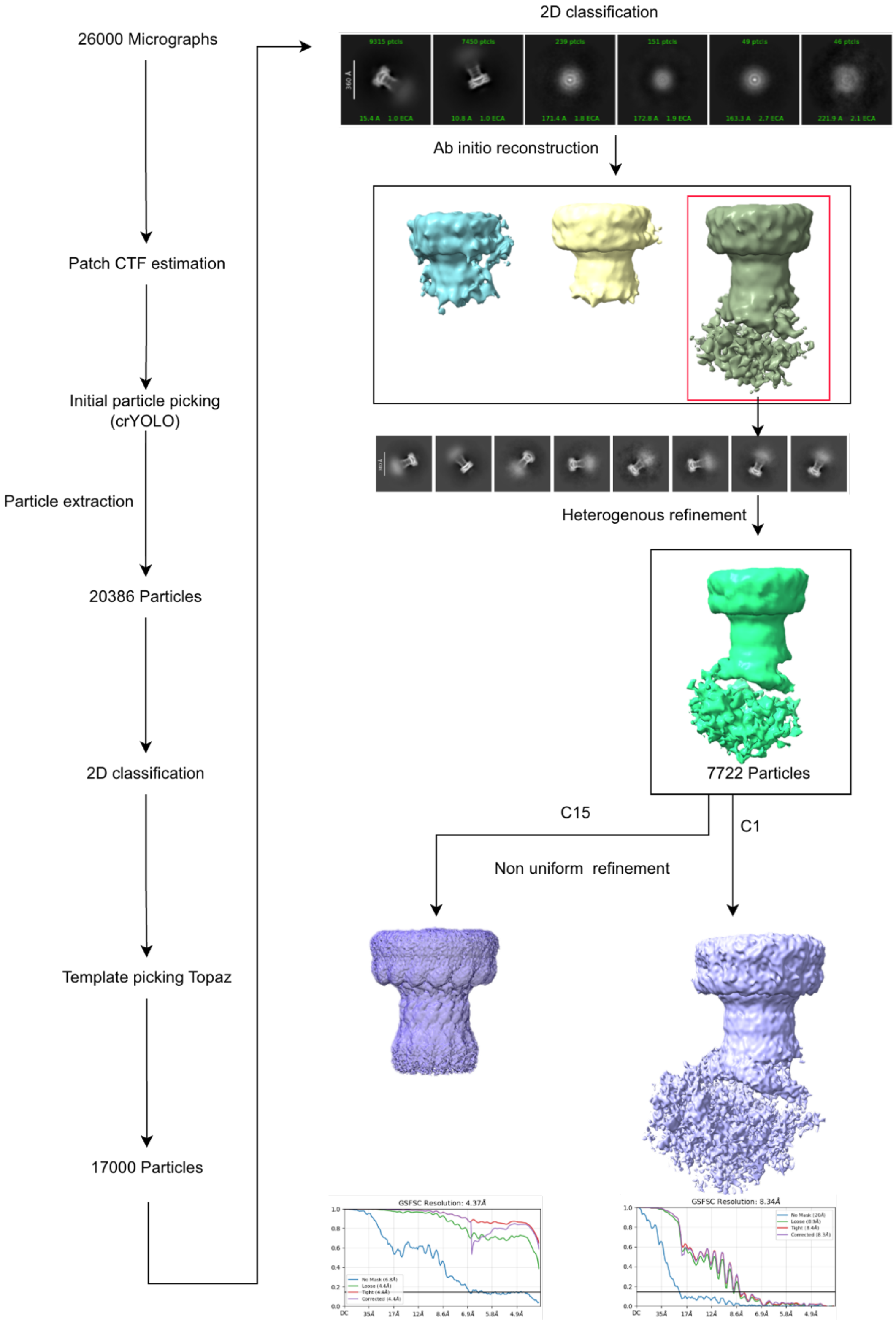


**Supplementary Figure 5. Cryo-EM single particle analysis for the *E.coli* IHE3034 type II secretion system.**

| Construct | Use | Method & primers |
|---|---|---|
| *pASK-IBA3C-gspOSCDEFGHIJKLM-Hui*<br><br>(O-M) | Initial construct to express the full T2SS | The primers used to clone the T2SS from *E. coli* O18:K1:H7 (IHE3034) genomic DNA. A double restriction digest of the *pASK* plasmid was used (restriction sites underlined).<br>F (*SacI*):<br>TCCGAGCTCCTTTTTGATGTTTTTCAGCAATACCCCGCGGCGATGCCCATACT<br>R (*XhoI*):<br>CCGCTCGAGGTGATGATGATGATGATG CCCCCGTCCAAACTCCAGCCGCTGCACATTCAC<br>Infusion was subsequently used to introduce the N-terminal Strep tag on GspE.<br><br>F: CCGCAGTTCGAAAAAGTGCCTGTAGCACAGGAAACCACCGCTAACACCGTGCGTCTGCCCT<br>R: TTTTTCGAACTGCGGGTGGCTCCACATTAACGCGTTCTCCCGGCATTGAGGAACGCGC |
| *pASK-IBA3C-gspE-gspLcyto*<br><br>(E/Lcyto) | Determination of the structure the ATPase | Infusion used for cloning.<br>The genes for GspE and GspL were cloned from genomic DNA into a single *pASK-IBA3C* vector to produce GspE with C-terminal His-tag. The cytoplasmic domain of GspL plus 21 transmembrane residues expressed.<br><br>Primers:<br>*gspE-his-vector*<br>F: GCGCTGGTAGTGGAAAGCGCTTGGAGCCACCCGCAGTTCGAAAAATAA<br>R: CAGCATCATAACCGTTAGTGATGATGATGATGATGCGCCTCCAT<br><br>*gspLcyto*<br>F:<br>CGGTTATGATGCTGGTGACCATCACGCTTTCCGGGCGGATGCAGCAACAACTTGGGCGAAC<br>R:<br>TTCCACTACCAGCGCAACCAGAATCAGCAATATCGGCAGAATCATCACCCGC |
| *pASK-IBA3C-gspEFGHIJKLM*<br><br>(E-M) | Demonstrates importance GspC in ordering the inner membrane assembly complex | Infusion used for cloning.<br>*pASK gspE-his-strep-gspM*. *BsaI* site of *pASK-IBA3C* used.<br>Primers:<br>F:<br>CGAGGGCAAAAAATGGTGCCTGTAGCACAGGAAACCACCGCTAACACCGTG<br>R:<br>GTGGCTCCAAGCGCTCCCCCGTCCAAACTCCAGCCGCTGCACATTCACCATCCCA |
| *pRSFDuet-sslE*<br><br><br><br>Co-transformed | Secretion of SslE by recombinant T2SS demonstrated | Double restriction enzymes followed by ligation used:<br>F(*NdeI*):GGAATTCCATATGAATAAGAAATTTAAATATAAG<br><br>R(X*hoI*):CCGCTCGAGTTACTTGTCGTCATCGTCTTTGTAGTCCTCGGCAGACATCTTATGC |

| | | |
|---|---|---|
| with (see above): *pASK-IBA3C-gspOSCDEFGHIJKLM* | | |
| *pRSFDuet-gspG*<br><br>Co-transformed with (see above): *pASK-IBA3C-gspOSCDEFGHIJKLM* | Pilus generation by T2SS demonstrated | Double restriction enzymes followed by ligation used.<br>F:(*BspHI*):CTAGTCATGAATTCGTTATCCCGCACACAAAAACCAC<br>R:(*BamHI)*:CGCGGATCCTTACTTGTCGTCATCGTCTTTGTAGTCCTGAAACTCCTGCAAATTCCAGTTACCGATATC |
| *GspOSCDEFGHIJKLM-Nouran* | Determination of CLM heterotrimer structure | A Flag tag was fused to the C-terminus of GspC using the *pASK-IBA3C-gspOSCDEFGHIJKLM-Hui* template and the following primer pair:<br>F:TCCATCGCACTGCGCGACTACAAAGACCATGAC<br>R:GATTAAATGCGGTTACTTGTCATCGTCATCCTTG, and<br>F:GATGACGATGACAAGTAACCGCATTTAATCCAG<br>R:CATGGTCTTTGTAGTCGCGCAGTGCGATGGAAATG<br><br>This template was then further adapted by GenScript Biotech (UK) to introduce a mini-GFP at the C-terminus of GspC and hexahistatidine tag at the C-terminus of GspE. |
| *GspOSCDEFGHIJKLM-Minoo* | Determination of secretin-pilotin complex<br><br>Determination of secretin-pilotin complex with HR<br><br>Determination of full secretion system GspSCDEFGHIJKLM | A Flag tag on the C-terminal of GspF wasintroduced into the *pASK-IBA3C-gspOSCDEFGHIJKLM-Hui* template.<br><br>Fflag-3034-F- GAT GAT GAT AAA TAA TTT ACG GAG TTA TCA CAT G<br>Fflag-3034-R- ATC TTT ATA ATC CAT TCC AAC CAT ATT GTT C |

**Supplementary Table 1. *E. coli* IHE 3034 constructs used this work.** Over 30 constructs were made in total; the seven above were used in work reported here.
